# Delineating the Transcriptional and Phenotypic Impact from Biotherapeutic Glycoengineering

**DOI:** 10.64898/2026.04.21.719832

**Authors:** Haining Li, Austin W.T. Chiang, Vahid H. Gazestani, Bokan Bao, Shangzhong Li, Patrice Ménard, Johnny Arnsdorf, Zulfiya Sukhova, Sara Petersen Bjørn, Karen Kathrine Brøndum, Anders Holmgaard Hansen, Sanne Schoffelen, Bjørn G. Voldborg, Nathan E. Lewis

**Affiliations:** Department of Bioengineering, University of California, San Diego, La Jolla, CA 92093, USA; Department of Pediatrics, University of California, San Diego, La Jolla, CA 92093, USA; The Novo Nordisk Foundation Center for Biosustainability at the University of California, San Diego, La Jolla, CA 92093, USA; Bioinformatics and Systems Biology Graduate Program, University of California, San Diego, La Jolla, CA 92093, USA; The Novo Nordisk Foundation Center for Biosustainability, Technical University of Denmark, Kongens Lyngby, Denmark; National Biologics Facility, Technical University of Denmark, Kongens Lyngby, Denmark; Dept. of Biotechnology and Biomedicine, Technical University of Denmark, Kongens Lyngby, Denmark; Complex Carbohydrate Research Center, Center for Molecular Medicine, and Department of Biochemistry and Molecular Biology, University of Georgia, Athens, GA

**Keywords:** Glycoengineering, N-linked glycosylation, Therapeutic protein, DNA damage and repair, Systems glycobiology

## Abstract

Glycosylation is critical to biopharmaceutical activity, stability, and pharmacokinetics. While production cells can be engineered to produce better protein glycoforms, glycans decorate thousands of host cell proteins, and it remains unclear how glycoengineering impacts the host cell. To decipher the cell response to glycoengineering, we studied a library of 166 glycoengineered CHO-K1 cell clones representing 54 different glycosyltransferase modifications. Through integrated analysis of glycomics, RNA-Seq, and phenotypic data, we discovered that glycoengineered mutants clustered into three distinct groups (wild-type-like, Moderate, and Substantial) based on their glycosylation patterns. Different glycosyltransferase families exhibited distinct phenotypic signatures: St3gal modifications increased growth rate and cell density, B4galt knockouts affected cell size, and Mgat knockouts enhanced cell viability. Notably, we found specific cellular reprogramming associated with each glycosyltransferase family, including alterations in energy metabolism, stress responses, and DNA repair mechanisms. These findings were validated in an independent set of 30 glycoengineered CHO-S cell lines, expressing a panel of 10 recombinant proteins. Our extensive analysis reveals phenotypic changes resulting from glycoengineering, identifies their molecular bases, and provides crucial insights for controlling glycosylation during therapeutic protein production.

## Introduction

Therapeutic glycoproteins are widely used for diverse diseases and disorders^1–4^. To enhance many of these drugs, glycoengineering has been used to modulate their activity, half-life, and immunogenicity^5–10^. Studies in animal models and human diseases have shown that changes in glycosylation machinery can have profound physiological effects, with glycosyltransferase deficiencies leading to various pathological conditions^11^, and many genetic variants likely impact proximal glycan structure^12^.Their significance is validated by regulatory agencies; the U.S. Food and Drug Administration (FDA) and the European Medicines Agency (EMA) require strict control of glycoprotein quality attributes due to glycans’ crucial role in therapeutic efficacy and half-life^13–17^.

Glycosylation can be controlled in host cells through various strategies (e.g., media optimization, chemical treatments, or genetic modifications^18–20^), and studies have successfully engineered CHO cells to produce more homogeneous N-glycosylation^21–23^. However, it remains unclear how such interventions impact the host cell. This knowledge gap is particularly concerning since glycans are involved in cellular processes including protein folding, trafficking, cell signaling, and cell-cell interactions. Understanding these effects is crucial not only for optimizing production but also for ensuring the stability and reliability of production cell lines.

The connection of glycosylation to host cell physiology is complex. Thus, valuable information can be obtained through the integration of multiple data types, shedding light on the underlying biological mechanisms^24^. Glycomics and phenomics provide essential data, reporting glycan structures^25^ and phenotypic readouts^26–28^, and the integration of additional omics, e.g., transcriptomics, provides valuable information to bridge the gap between glycotype and phenotype^29^. Specifically, quantifying the expression of genes associated with adding, removing, remodeling, and transporting sugar molecules can suggest the mechanisms underlying the repertoire of glycans observed^30,31^, while analysis of the rest of the transcriptome can shed light on other pathways and processes connected to phenotype^29^. This multi-omics integration is particularly useful as it can reveal molecular bases for phenotypic changes and identify compensatory mechanisms in the glycosylation machinery that might be missed by analyzing any single data type in isolation^24^.

Here, we study the impact of glycoengineering on a library of 166 glycoengineered CHO-K1 (geCHO-K1) cell lines, representing 54 different combinations of glycosyltransferase knockouts and knockins (Table S1). Through an integrated multi-omics analysis of glycomics, transcriptomics, and phenotypic profiling across these geCHO-K1 cell lines, we investigated three fundamental questions: 1) How do genetic modifications of specific glycosyltransferases affect the biosynthetic steps in glycosylation? 2) How does the knockout of different glycosyltransferases impact cellular phenotypes? and 3) Which metabolic functions and pathways are altered when glycosyltransferases are knocked out or in? Our analysis revealed that mutants cluster into three distinct groups based on their glycan synthetic patterns compared to wild type, with each group showing unique phenotypic signatures. Notably, we discovered that different glycosyltransferase families (Mgat, B4galt, St3gal) exhibited distinct correlations with phenotypic and pathway alterations. For instance, perturbations in the St3gal family led to considerable changes in metabolic pathways, ion transport, and DNA repair processes. To ensure robustness of our findings, we validated the major phenotypic perturbations through analysis of an independent set of 29 glycoengineered CHO-S clones (geCHO-S) producing 10 different recombinant proteins.

Through this systematic analysis, we illuminate the complex interplay between glycoengineering and cellular homeostasis, revealing pathways associated with glycosylation changes and phenotypic alterations. Our findings provide critical insights into the broader cellular impact of glycoengineering and underscore the fundamental importance of glycosylation in mammalian cell biology. These insights have important implications for optimizing biopharmaceutical development and manufacturing processes.

## Results

### A panel of 166 clones with 54 different glycoengineering designs

To investigate the cellular impacts of altered glycosylation, an extensive panel of glycoengineered CHO cell lines was developed and characterized through multiple stages. The study design consisted of five major phases (Fig. 1). Initially, a panel of glycoengineered CHO-K1 cell lines using zinc-finger nucleases (ZFNs) to create targeted knockouts of glycosyltransferase genes^21^. These engineered lines were then used to produce and analyze recombinant erythropoietin (EPO), with glycoprofile data from their distinct genotypes reported previously^21^.

**Figure 1.**
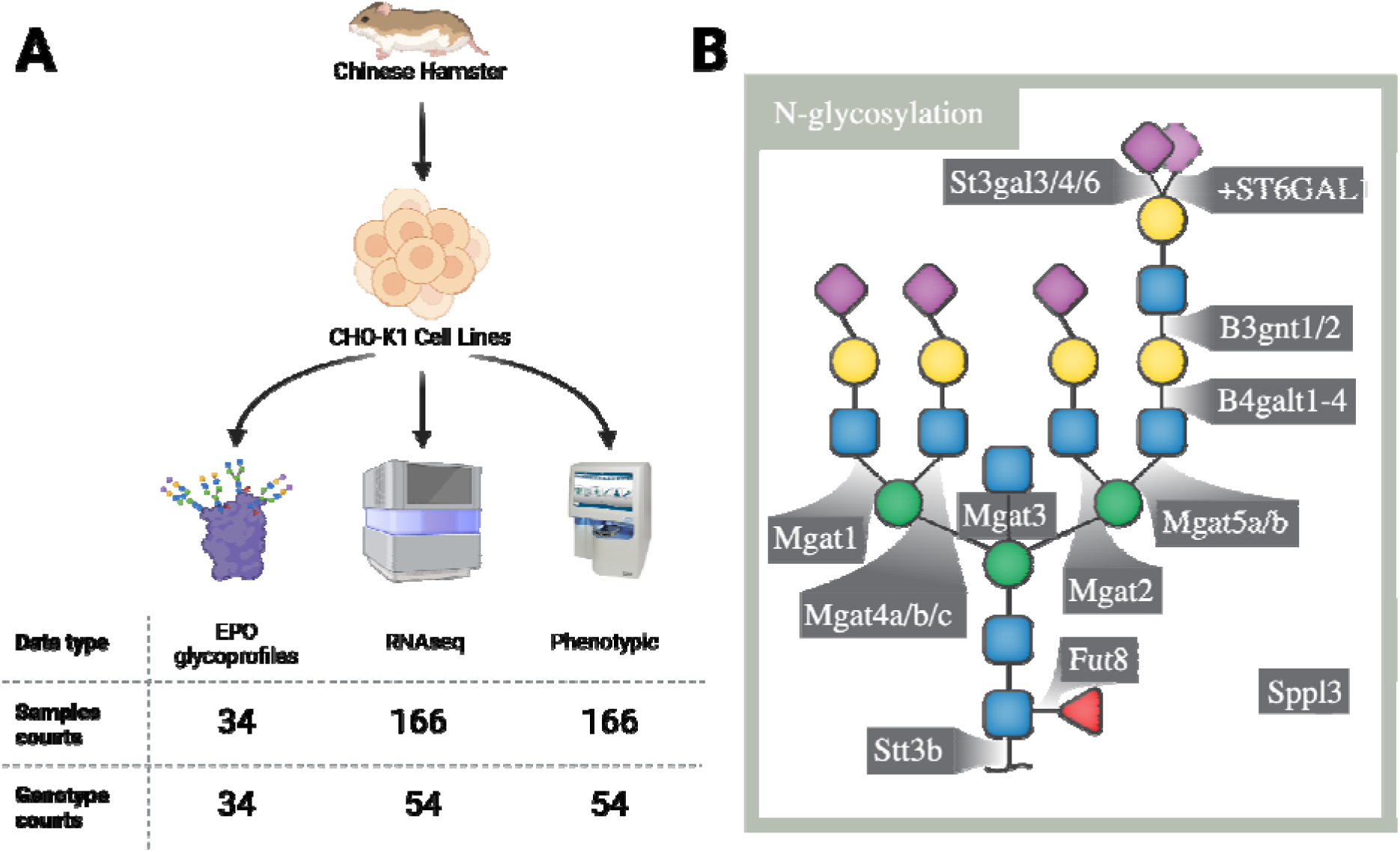
Experimental workflow for characterizing glycoengineered CHO (geCHO-K1) cell lines. **A)** The data were generated from CHO-K1 cells with combinations of glycosyltransferases knocked out or knocked in. We analyzed previously-published^21^ EPO glycoprofiles from 34 of the genotypes. Then, we performed an analysis of an expanded panel with 54 genotypes (166 total samples), including RNA-Seq and phenomic characterization. **B)** Key glycosylation-related enzymes targeted in this study, including genes involved in N-glycan branching (Mgat1, Mgat3, Mgat4a/b/c, Mgat5a/b), sialylation (St3gal3/4/6, St6GAL1), galactosylation (B4galt1-4), fucosylation (Fut8), and other glycan-modifying enzymes (B3gnt1/2, Stt3b, Spp13). Knockouts (e.g., St3gal3/4/6) and knock-ins (e.g., +ST6GAL1) were designed to perturb specific glycan biosynthesis pathways, enabling systematic analysis of their roles in glycosylation efficiency and product quality.

Building on this foundation, we expanded the characterization of these cell lines through comprehensive molecular and phenotypic profiling. From the original panel, we obtained 54 unique genotypes, each represented by approximately 3 biological replicates, yielding 166 total samples. These samples were subjected to RNA-Seq, and phenotypes were assayed with the Bioprofile FLEX Analyzer (measures key cellular phenotypes including cell size, growth rate, viability, and concentrations of glucose, lactate, glutamine and glutamate). The multi-omics dataset was then analyzed to identify molecular signatures and pathway alterations associated with specific glycoengineering modifications.

To validate our findings, we generated an independent panel of 29 geCHO-S cell lines derived from the CHO-S lineage, and studied these as they transiently expressed 10 different recombinant proteins. This validation cohort was subjected to RNA-Seq and glycoprofiling of the recombinant proteins of interest to confirm the key molecular and glycoform changes observed in the primary CHO-K1 panel.

### Dominant isozymes and off-target effects are apparent from glycoprofiles

Glycosyltransferases are vital for eukaryotic physiology, with enzyme deficiencies potentially altering cell functions^32–34^. However, to understand how glycoengineering impacts the host cell, we first need to assess which genetic changes induce a meaningful change to glycosylation. EPO was previously expressed in 34 different glycoengineered cell line genotypes and subsequently glycoprofiled^21^. We reanalyzed these data using Glycompare^35^ to quantify all glycan intermediates (glyco-motifs) for each of the 34 EPO glycoprofiles^21^. This allowed us to identify glycoengineering designs that led to structurally similar glycans. After obtaining the glyco-motif abundance profiles, we conducted hierarchical clustering to identify mutants with similar glycan biosynthesis.

Upon analyzing the EPO glyco-motifs, we found the mutant cell lines clustered into a few groups with similar glycoprofiles (**Figure 2**). In particular, several single-glycosyltransferase knockouts clustered tightly with the wild-type parental CHO-K1 cell line. This “Wild type-like” group (green dendrogram), have mutations in glycosyltransferases that are either minimally functional, are supported by active and redundant isozymes. The next cluster, or “Moderate” group (blue dendrogram), included genotypes with 1 or 2 glycosyltransferase knockouts. As expected, with additional knockouts, we observed a clear clustering of samples with extensive changes in glycosylation, which we designate as the “Substantial” group (red dendrogram).

**Figure 2.**
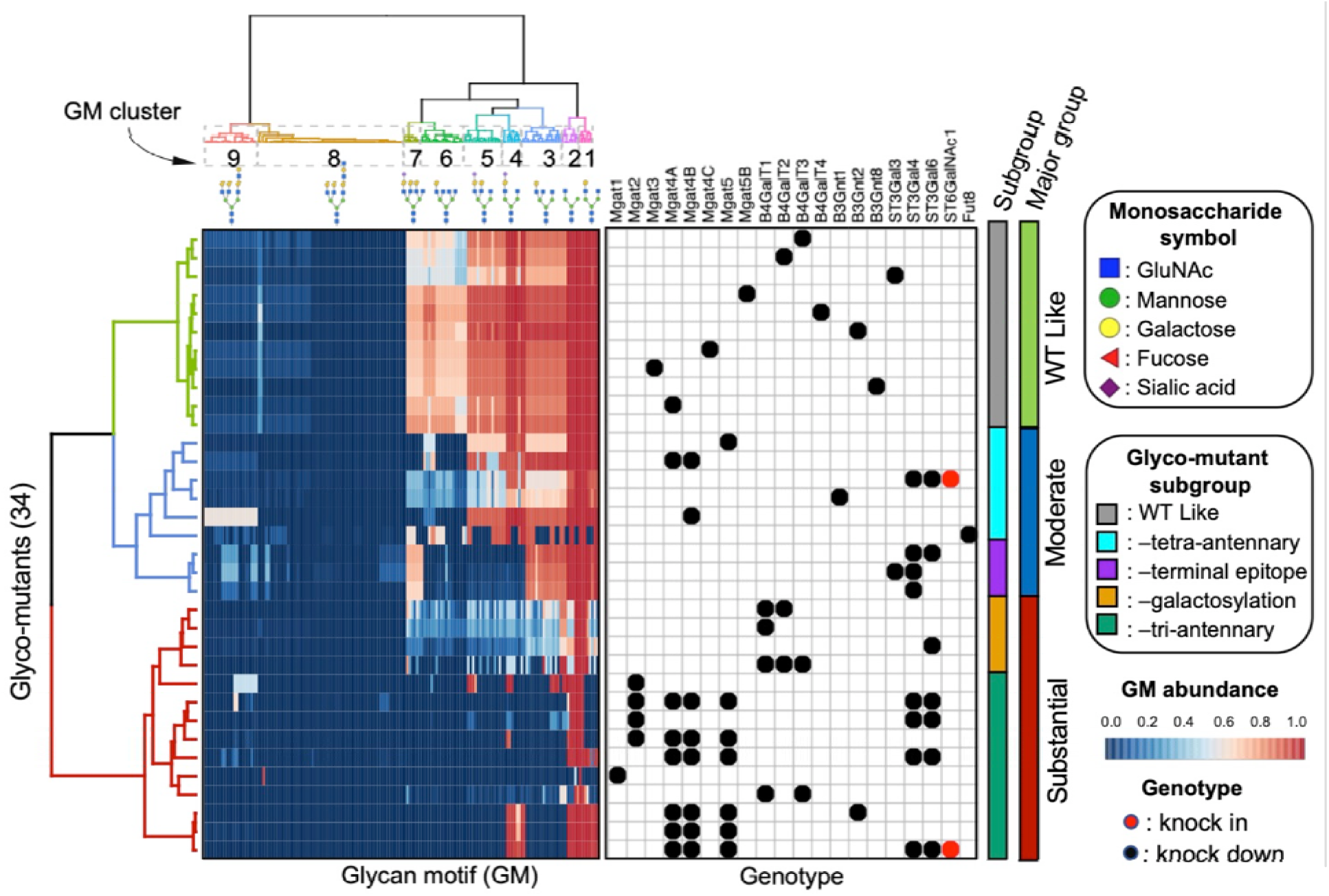
Glycoprofiles of geCHO-K1-produced EPO clustered into products from cells with similar genotypes. Glycoprofiles were processed using GlyCompare and clustered to identify mutants with similar glycoprofiles. Clustering was driven by nine glyco-motif clusters (GM 01-09), in which each cluster is annotated with their most representative glycan topology (greater than 60% of the glyco-motif member within a cluster share the same substructure). There are three primary clusters of mutants on the 34 glycoengineered samples, named by the degree of perturbation: WT-like (green), Moderate (blue), and Substantial (red).

Each group included knockouts involved in similar glycan synthesis pathways. As expected, clones with multiple sialyltransferases targeted (e.g., St3gal4/6, St3gal3/4, and St3gal4) grouped together, separate from the ‘Wild type-like’ group. Similarly, samples from clones wherein Mgat5, Mgat4a/4b, and Mgat4b were knocked out, also clustered more tightly and showed a decrease in the tetra-antennary-related glyco-motif clusters (GM clusters: GM6 and GM7).

The clustering of glyco-motifs informs which glycosyltransferases cause minor modifications and which combinations disrupt the wild-type glycoform. Indeed, different isozymes within the same glycosyltransferase family could show different magnitudes of impact on the glycoprofiles. For example, we knocked out multiple St3gal family isozymes (i.e., St3gal3, St3gal4, and St3gal6), but only mutants with St3gal4 knocked out showed a decrease in sialylation; meanwhile, the St3gal3 mutant showed “wild type-like” glycoprofiles and the St3gal6 knockout clustered as “Substantial”, due this clone resulting in decreased lacNAc elongation compared to the other St3gal KOs. Similar observations were found with the Mgat family. Dominant isozymes in the moderate and substantial groups included Mgat4b/5 and B4galt1, St3gal4, and St3gal6.

### Changes in glycosylation co-varies with compensatory glycosyltransferase gene expression

Glycoprofile clustering identified mutants with similar glycosylation. However, do the cell lines show compensatory gene expression? We next tested if the changes in glycosylation result in differential gene expression of the remaining glycosyltransferases. Here, we first tested if unexpected changes in the glycome (i.e., beyond the expected changes from the known enzyme knockouts) could be explained by further transcriptomic changes. This survey identified a short list of genes with variations sufficient for delineating a transcriptomic signature specific for each glycoengineered mutant (**Figure 3**). We compared the clustering of the mutant glycoprofiles and the transcriptomic signature for each mutant, and found that both data types clustered similarly (**Figure 3A**), as quantified using the Fowlkes-Mallows score (FMS) (p = 2 x 10^-4^) (**Figure 3B**). The clusters have an FM index of 0.64, indicating a high level of agreement between the glycosylation clusters and the gene expression clusters.

**Figure 3.**
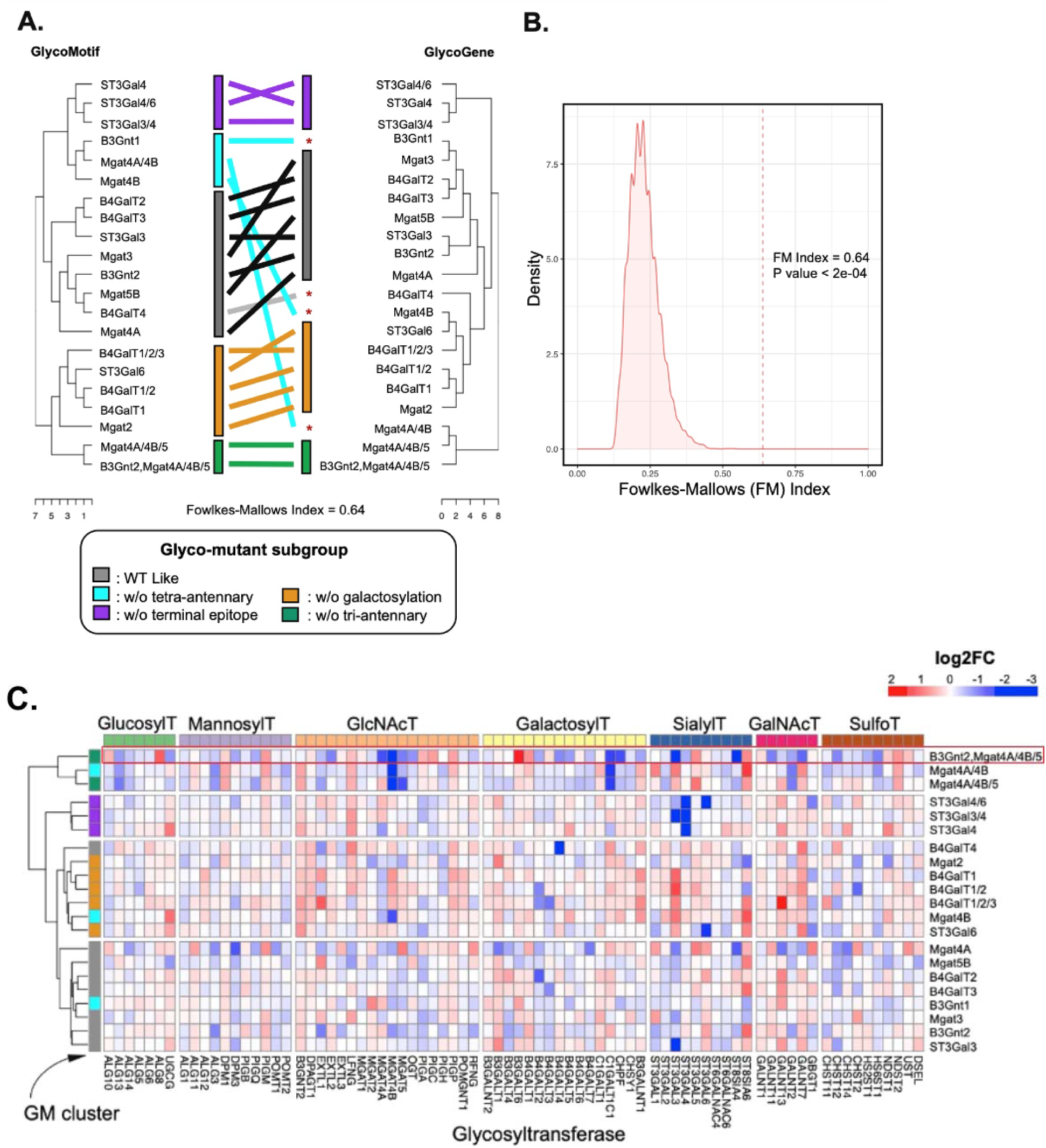
Non-targeted glycosylation machinery can be impacted by glycoengineering. **A)** Glycoprofiles and gene expression of mutants clustered similarly. **B)** The FM index suggests a high similarity between the two clusterings. The distribution of randomized FMS across various clustering comparisons is visualized in a density plot, with the FM index plotted along the x-axis and the density values along the y-axis **C)** Differential gene expression analysis shows changes in the expression of additional glycosyltransferases beyond the target genes.

A subsequent differential gene expression analysis (**Figure 3C**) showed multiple glycosyltransferase genes were impacted in some mutants, beyond just the targeted genes. For example, in the mutant “B3gnt2/Mgat4a/4b/5”, St3gal3 and St8sia4 were down-regulated, and B3galt6 was up-regulated (highlighted in **Figure 3C**). Taken together, we conclude that glycosylation changes lead to gene expression changes in the glycosylation machinery.

### Stratification of phenotype induced by glycoengineering

A common concern in engineered cells is how they might impact cell phenotype or host cell survival. To explore the phenotypic relationships among the geCHO-K1 mutants, we analyzed the phenotypic profiles of the 54 geCHO-K1 mutant genotypes using the Bioprofile FLEX analyzer. Hierarchical clustering showed that many of the sialyltransferase (e.g., St3gal3, St3gal4, St3gal6) knockout mutants clustered tightly as a group (cluster B01 in **Figure 4A, green dendrogram**), while other mutants (e.g., B4galt1, B4galt2, B4galt3, B4galt4) made a second cluster with modest increases in cell size and osmolarity. A third cluster, B03, included many mutants targeting branching-related glycosyltransferases (e.g., Mgat4a, Mgat4b, and Mgat5); these showed greater increases in viability, lactate, and ammonium concentrations.

**Figure 4.**
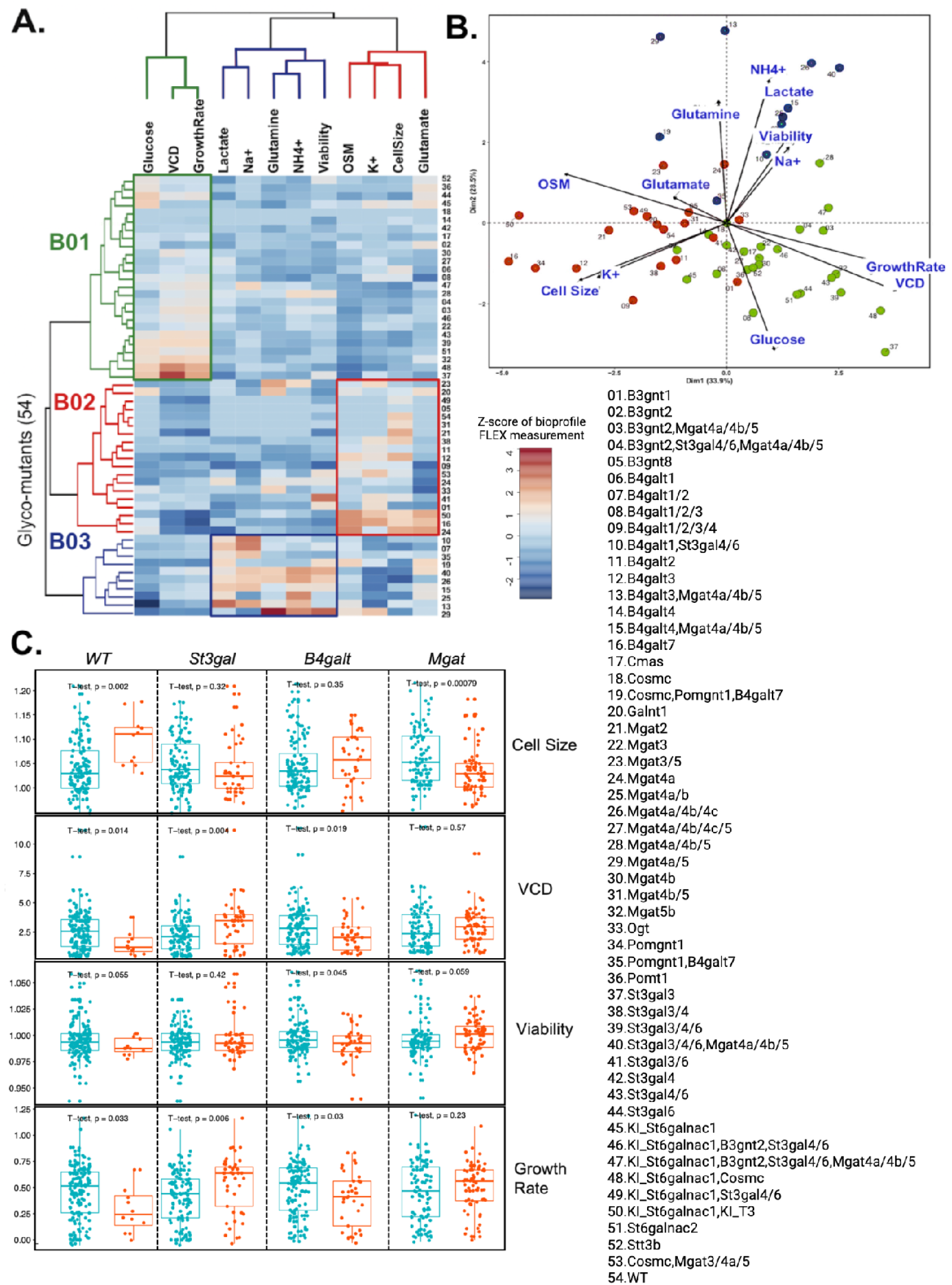
Bioprofile FLEX phenotype clustering. (A) There are three dominant row clusters on the 54 glycoengineered samples: B01 (Growth rate and VCD; green), B02 (Cell size; red), and B03 (Viability; blue). (B) Principal component analysis on the 54 glycoengineered samples. Dots are colored to correspond to the cluster colors in (A). (C) Phenotypic alterations in different glycosyltransferase-family knockout.

To further explore the clustering of these phenotypic variations, we performed Principal Component Analysis (PCA) (**Figure 4B**). The first PC accounts for 33.9% of the total variation and separates the glycoengineered mutants with phenotypes by Cell Size and Growth Rate. The second PC accounts for 28.5% of the variation and separates the cell lines predominantly by viability. When we visualized the samples according to hierarchical cluster assignments from Figure 4A, the PCA visualization revealed major phenotypic features of the three clusters (**Figure 4B**).

To more clearly associate mutant genotypes with phenotypes, we conducted targeted analyses to ascertain which glycosyltransferase mutations exert the most pronounced effects on specific phenotypic features (**Figure 4C**). We focused on the knockout effects of St3galt, B4gal, and Mgat families on four distinct phenotypic features: cell size, VCD, viability, and growth rate. Among these comparisons, we found notable changes in Cell Size following the knockout of Mgat (t-test, p-value = 0.0008); VCD and Growth Rate following the knockout of St3gal (t-test, p-value = 0.004 and 0.006, respectively); and Viability, VCD, and Growth Rate following the knockout of B4galt (t-test, p-value = 0.045, 0.019 and 0.03, respectively). These findings show relationships between particular glycosyltransferase mutations and their phenotypic manifestations.

We next integrated phenotypic and gene expression data using Similarity Network Fusion (SNF)^36^ to reveal molecular mechanisms associated with cellular response to glycoengineering. SNF allows one to combine distinct data types into a unified similarity network of samples (see Methods). The sample groups represent subsets of the most similar cellular responses and gene expression profiles. SNF clustering on the integrated data revealed three latent molecular phenotype (MP) subgroups in the geCHO-K1 cells (**Figure 5**), indicating differences in the cellular responses to different glycoengineering strategies, and each subgroup exhibits distinct transcriptional signatures. The identified subtypes are strongly associated with the PCA clusters (**Figure 4B**). Specifically, the ‘MP01’ mutants (red) exhibited significantly elevated ‘Cell size’ but reduced ‘Growth rate’ and ‘Viability,’ and the ‘MP02’ mutants (green) exhibited significantly increased ‘Ammonia’ and ‘Lactate.’ Interestingly, the ‘MP03’ mutants (blue) showed an opposite phenotype to the ‘MP01’ mutants, which significantly enhanced ‘Growth Rate’ and ‘Glucose consumption rate.’ To characterize the dysregulated pathways associated with the variability of cellular responses exhibited by the identified MPs, we assessed the importance of all Reactome^37^ pathways. Our results show that ‘MP01’ is associated with PC3 and PC4 pathways, ‘MP02’ is associated with PC1,2,3,5 and 7 pathways, and ‘MP03’ is associated with PC6 pathways.

**Figure 5.**
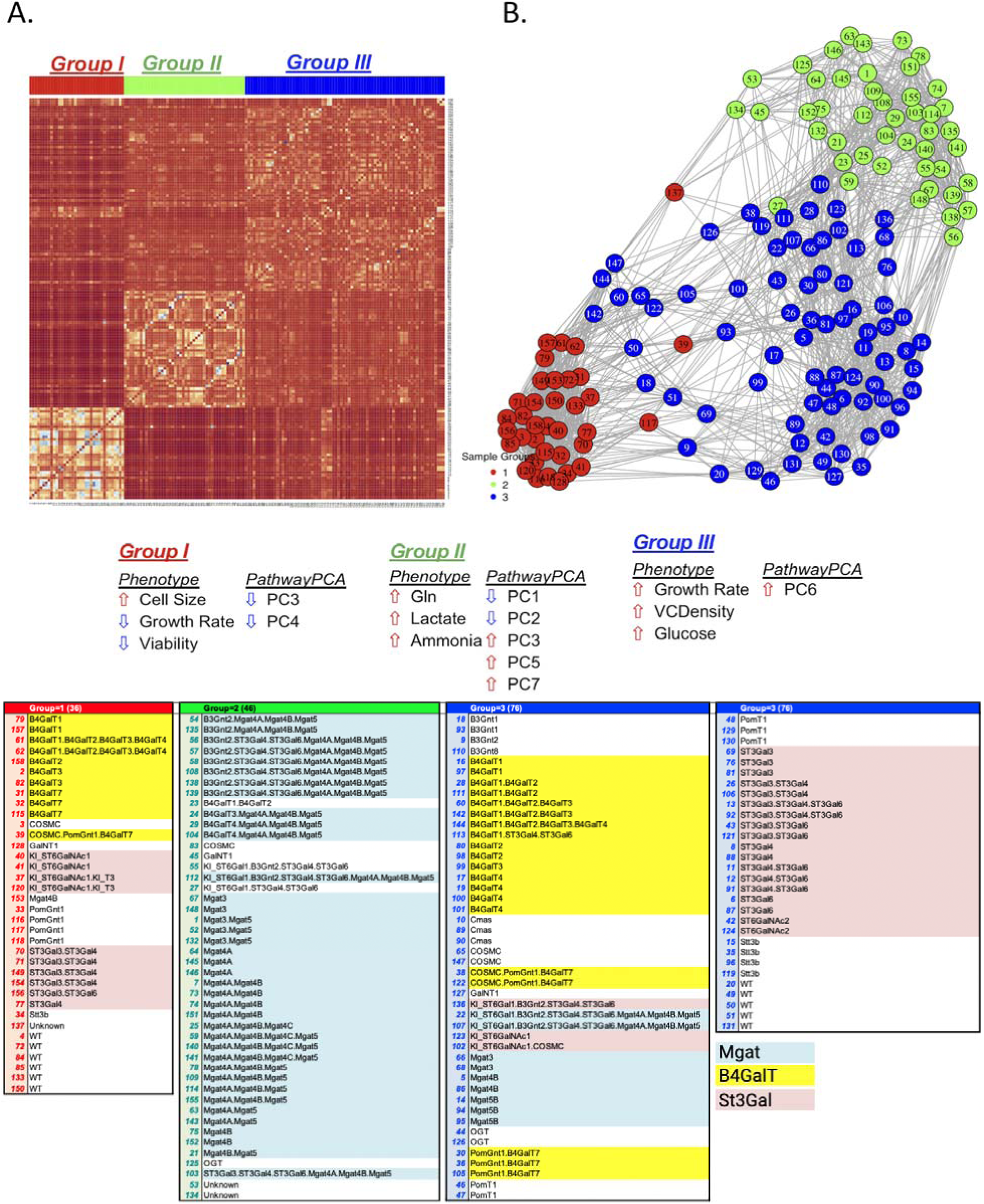
Pathway and phenotypic variation clustered glycoengineered mutants into three groups based on Similarity Network Fusion (SNF). (A) SNF was conducted on the transcriptomic and phenotyping data of the 166 geCHO-K1 cells and (B) SNF clusters were visualized, and groups were evaluated for their predominant phenotypic and transcriptional signatures.

### Metabolic reprogramming underlying cellular response induced by glycoengineering

CHO cell metabolism is a major driver of cell productivity and product quality^38–42^. To determine if the geCHO-K1 cells exhibit significant metabolic changes contributing to the altered phenotypes, we performed a gene set variation analysis (GSVA) to identify significant alterations in phenotypic traits. Utilizing the GSVA results, we then conducted ordinal regression to identify biological pathways that were increasingly dysregulated with increasing numbers (0-3) of isozymes targeted in mutant cell lines. Below, we summarize the findings across different glycosyltransferase families.

The Mgat-family mutants exhibit significant changes in many pathways, and here we present the top 3 up- and 3 down-regulated pathways that increase or decrease with the knockout of increasing numbers of Mgat-family genes. Notably, we found increased activities in pathways involving activated point mutants of FGFR2, regulation of the pyruvate dehydrogenase (PDH) complex, and regulation of Rheb GTPase activity by AMPK (**Figure S2B, a-c**). The activation of FGFR2 suggests an enhancement in oncogenic signaling, potentially promoting cell proliferation and survival^43^. The increased activity of PDH complex regulation likely reflects cellular adaptation to handle elevated pyruvate levels resulting from increased glycolysis^44^. Furthermore, upregulation of AMPK activity and subsequent inhibition of Rheb GTPase and mTORC1 signaling can be a compensatory response to metabolic stress, possibly through enhanced autophagy to maintain cellular homeostasis^45,46^. These pathway activities, combined with our observation of increased lactate secretion, suggest a metabolic reprogramming toward Warburg metabolism^47^. This metabolic phenotype is commonly observed in rapidly proliferating cells^48^, and inhibits cell growth and productivity in CHO cells^40–42,49,50^. Conversely, pathways were downregulated associated with p75NTR signaling complexes, acyl-chain remodeling of phosphatidylinositol (PI), and SHC1 events in ERBB4 signaling (**Figure S2B, d-f**). The downregulation of p75NTR signaling may promote survival and inflammatory responses, potentially reducing apoptosis^51^. Reduced acyl-chain remodeling of PI could affect PI’s role in regulating Warburg metabolism through the PI3K/AKT/mTOR pathway^52^. These results were consistent with our observation of increased lactic acid production (**Figure 5**). Finally, the decreased activity in SHC1-mediated ERBB4 signaling suggests a reduction in proliferative and survival signals, which could impact overall cell growth^53^.

The B4galt-family mutants, on the other hand, exhibit enhanced stress responses and reduced critical signaling pathways with increasing the number of B4galt genes knocked out. We observed an upregulation in the pathway activities involving acyl-chain remodeling of phosphatidylglycerol (PG), the unfolded protein response (UPR), and the calnexin-calreticulin cycle (**Figure S3B, a-c**). The upregulation of acyl-chain remodeling suggests enhanced lipid metabolism and membrane dynamics, likely as an adaptive response to maintain membrane integrity under stress conditions^37,54^. The increased activity in the UPR indicates substantial ER stress as cells attempt to cope with the accumulation of misfolded proteins, a critical process for maintaining cellular homeostasis^55,56^. Furthermore, the upregulation of the calnexin-calreticulin cycle could impact protein folding and processing^57,58^ (**Figure S3B, d-f**).

The dominant pathway changes that correlated with the number of mutated St3gal genes included those involved in metabolism, ion transport, and DNA repair. When more St3gal isozymes were knocked out, the Endogenous Sterols and Reversible Hydration of Carbon Dioxide pathways were upregulated (**Figure 6B, a-c**). The Endogenous Sterols pathway involves the biosynthesis and regulation of sterols, which are essential components of cellular membranes and cell-cell communication^38^. The Reversible Hydration of Carbon Dioxide pathway is crucial also for maintaining acid-base homeostasis through the reversible conversion of carbon dioxide and water to bicarbonate and protons^37^. Downregulated pathways included Homologous Recombination Repair of Replication Independent Double Strand Breaks, and Double Strand Break Repair (**Figure 6B, d-f**). This could suggest more compromised genomic stability, increasing the risk of mutations, chromosomal aberrations, and production instability^59–61^.

**Figure 6.**
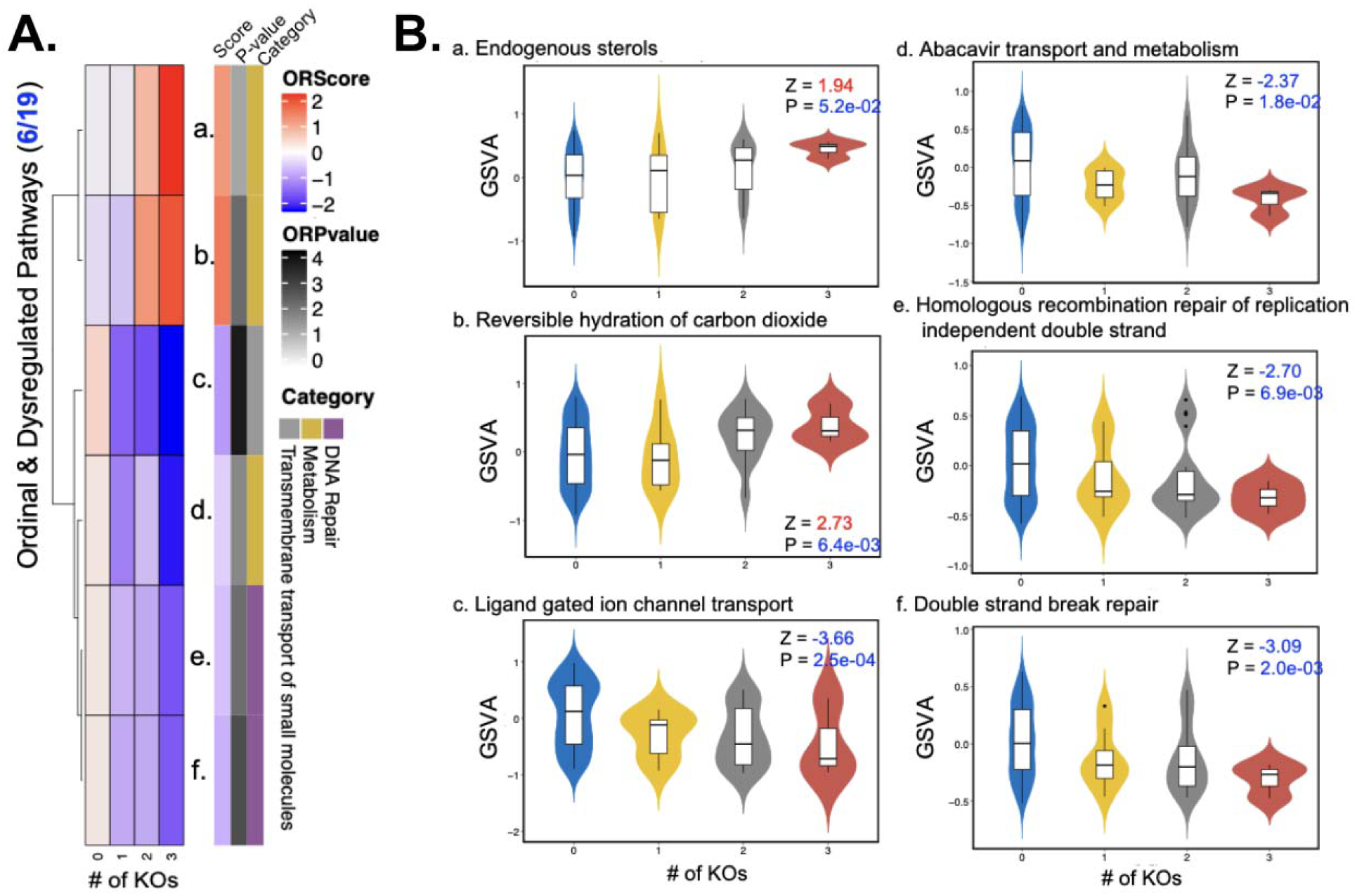
Ordinal regression results of GSVA of pathways in the St3gal-family glycoengineered strains. (A) The heatmap of statistically significant dysregulated pathways with an ordinal trend of GT knockouts. (B) Statistically significant dysregulated pathways.

To corroborate our findings of St3gal gene mutations impacting expression of DNA repair machinery, we performed similar analyses using an independent dataset of 363 glycoengineered CHO-S (geCHO-S) samples. In this dataset, there are 29 geCHO cell lines comprising different knockout/in combinations of 17 glycosyltransferase gene combinations to produce a range of different glycoforms of ten recombinant glycoproteins, including monoclonal antibodies, fusion proteins, cytokines, and enzymes. We performed RNA-Seq to assess the transcriptome of this dataset. We tested if DNA-damage-repair-related pathways exhibited significant changes in expression with increasing KOs of St3gal genes. Despite being an independently-generated cell line panel, implemented in a different CHO cell lineage, we found that double-strand break repair is suppressed. This pathway showed significantly different enrichment scores between 0 knockouts (mean = 0.215) and 3 knockouts (mean = −0.299) (Mann-Whitney U test, p < 0.001) (**Figure S4**).

## Discussion

Here, we studied how CHO cells respond to glycoengineering. First, we found glycoprofiles from EPO produced in geCHO-K1 cell lines cluster into three distinct groups (wild-type-like, Moderate, and Substantial), with these glycomic changes strongly correlating with specific transcriptional signatures. Second, we discovered that perturbations to different glycosyltransferase families yield distinct cellular responses: St3gal modifications increased growth rate and cell density, B4galt affected cell size, and Mgat decreased cell size. Third, we observed metabolic reprogramming specific to different glycosyltransferase families, including alterations in energy metabolism in Mgat-family mutants, stress responses in B4galt KOs, and DNA repair mechanisms in St3gal mutants. These findings have important implications for production of glycoengineered biotherapeutics. The observed compensatory transcriptional responses, particularly in non-targeted glycosyltransferases, suggest that cells actively maintain glycan homeostasis through complex regulatory networks. Moreover, the perturbation of critical cellular pathways, such as DNA repair in St3gal mutants, indicates that glycoengineering can have broader impacts on cell physiology than previously appreciated. Understanding these cellular responses is crucial for developing more effective and stable glycoengineered cell lines for biotherapeutic production.

### The geCHO-K1 cells hint at interactions between glycan biosynthetic pathways

As expected, different glycoengineering approaches yield distinct glycoforms (**Figure 2**, left panels). However, compensatory mechanisms may exist within the complex glycosylation system to maintain glycan epitope homeostasis. Homeostasis is crucial for robustness in living systems. Several studies have reported examples illustrating how the glycosylation machinery maintains glycan epitope homeostasis^62,63^. Specifically, gene mutations in glycosyltransferases within the glycan biosynthesis pathway can result in the global compensation of missing glycan epitopes in other glycans. For example, Mgat4a and Mgat4b enable the synthesis of the GlcNAcβ1-4 branch on the Manα1-3 arm of the N-glycan core. This branch increases glycan epitope density in N-linked glycoproteins, enhancing overall lectin binding to N-linked glycans. Mgat4a/b double mutant mice exhibit significantly decreased N-glycan branches. However, the missing N-linked branch is observed as LacNAc epitopes on the remaining linear arms of the N-linked glycans^64^. These results demonstrate that the glycosylation machinery can effectively maintain glycan epitope density at homeostatic states. Although the underlying mechanisms by which the glycosylation machinery maintains glycan epitope homeostasis are mainly unknown, elucidating these mechanisms will undoubtedly reveal novel aspects of glycosylation control.

Our observations suggest that glycosylation alterations are influenced by several dominant isoenzymes (**Figure 2**). Specifically, the glycoform resulting from Mgat4b knockout exhibits a pronounced reduction in tetra-antennary glycans (GM cluster 6-7) compared to Mgat4A knockout’s glycoform. Moreover, B4galt1 knockout leads to a significant reduction in many hybrid glycans (GM cluster 1, 3-7) than knockouts of other isoenzymes: B4galt2, B4galt3, and B4galt4. Lastly, St3gal4 appears to be the dominant isoenzyme affecting the expression of tri-and tetra-antennary glycans (GM cluster 4-7), as also supported by a previous study^65^. These findings provide insights into how glycan compensatory mechanisms could be elucidated by the biochemical network.

### Metabolic shifts may compromise glycoengineered cells

Based on the observed pathway activity changes in the Mgat-family glycomutants, several conclusions can be drawn that relate to mutant-specific cell size and growth rate. The observed significant upregulation of pathways involving activated FGFR2, PDH complex regulation, and AMPK-mediated Rheb GTPase regulation suggests a metabolic shift toward the Warburg effect, which is characterized by high glycolytic activity even in the presence of oxygen. Supporting this metabolic shift, we observed increased lactate secretion as more genes were knocked out in Mgat mutants. This metabolic phenotype is commonly observed in rapidly proliferating cells, with its enhanced glycolysis for both energy production and biosynthetic needs. Conversely, we observed downregulation in pathways associated with acyl-chain remodeling of phosphatidylinositol (PI) and SHC1 events in ERBB4 signaling. Reduced acyl-chain remodeling of PI could affect membrane composition and metabolic signaling, potentially impacting lipid metabolism and glycolysis, thus impacting growth. The decreased activity in SHC1-mediated ERBB4 signaling suggests a reduction in proliferative and survival signals in these production cells. These pathway alterations align with our Bioprofile FLEX analysis, which showed smaller cell size and higher growth rate in Mgat mutant cell lines, though the precise mechanistic connections between these pathway changes and phenotypes require further investigation.

Our data suggests that B4galt-family mutants decreased in viable cell density (VCD) and growth rate. This was accompanied by a significant upregulation in pathways related to enhanced lipid metabolism, adaptive membrane dynamics, and substantial endoplasmic reticulum (ER) stress due to the accumulation of misfolded proteins. The associated stress can negatively impact cell growth and viability as cells prioritize survival and maintenance functions over proliferation. Future work should evaluate the performance of stable production clones of these cells to evaluate their susceptibility to ER stress.

### Glycoengineering can impact DNA damage repair and genome integrity

The Reactome pathway enrichment analysis following St3gal knockout revealed significant metabolic and activity pathway alterations (**Figure 5**). Notably, the upregulated pathways likely represent compensatory mechanisms to maintain cellular homeostasis under altered glycosylation conditions, which can support increased cell membrane integrity and efficient pH regulation, thereby promoting cell growth and survival. The upregulation of metabolic pathways might be an adaptive response to the stress induced by the knockouts, aiming to stabilize cellular functions. However, we also observed the downregulation of DNA repair mechanisms, potentially impacting genomic integrity^60^. This connection of suppressed sialylation to increased susceptibility to DNA damage is supported by multiple studies that have shown that the loss of sialylation can increase DNA damage vulnerability. For example, ST6GAL1 can protect against radiation-induced apoptosis in vitro^66,67^, and *in vivo* ST6GAL1 KO increased radiation-induced gastrointestinal damage^68^. ST6GAL1, has also been linked with resistance to human rectal cancer treatments, and its knockdown in organoid models increased chemoradiation-associated apoptosis ^69^. The reduced DNA repair efficiency suggests that the cells are prioritizing growth and proliferation at the potential cost of long-term genomic stability.

A critical challenge in biopharmaceutical development is establishing mammalian cell lines that can efficiently and effectively perform glycoengineering of therapeutic proteins^70–74^. While numerous cell lines have been developed, productivity loss remains a major concern in biomanufacturing. Research has demonstrated that this productivity instability often stems from genome instability^75,76^, where chromosomal abnormalities lead to reduced transgene copy numbers and decreased protein titer, as genomic instability disrupts the integrity of integrated transgenes^61^. Recent studies have revealed that deficiencies in DNA damage repair pathways can contribute to this genome instability^77,78^. Given these findings, investigating whether glycoengineered mutants exhibit significant alterations in DNA damage and repair mechanisms becomes crucial, as these changes could either benefit or harm product production and host cell survival. Understanding the underlying mechanisms of these cellular responses could provide valuable insights for rational optimization of expression systems, ultimately advancing the development of cell lines for sustainable glycoengineered therapeutic protein production.

## Conclusions

Here, we demonstrated that glycoengineering not only impacts glycosylation but that it yields phenotypic alterations in CHO cell lines. Furthermore, these molecular mechanisms are linked to the phenotypic alterations as glycosylation changes. Studies have shown that metabolism is associated with several CHO cell phenotypes^41,79,80^. Our findings provide insightful knowledge on the effects of glycoengineering and detailed mechanisms about how glycosylation and pathway interplay can lead to phenotype changes following glycoengineering on the CHO cells. Thus, these insights provide further tools to enable the development of a robust host with desired phenotypes when performing glycoengineering on the CHO cells for biotherapeutic products.

## Methods

### Glycoprofile analysis

Glycoprofiles of EPO, were already measured and published^21^. EPO glycoprofiling was performed on 34 different glycoengineered genotypes using MALDI-TOF (matrix-assisted laser desorption/ionization–time of flight) analysis of released permethylated N-glycans. For each sample, approximately 25 μg of purified EPO was digested overnight at 37°C with 2U PNGase F in 50 mM ammonium bicarbonate to release the N-glycans. Released N-glycans were separated using C18 Stagetips and incubated in 100 mM ammonium acetate (pH 5.0) for 60 minutes at 22°C.

The glycan samples were permethylated and extracted with chloroform. The organic phase was washed repeatedly with deionized water for desalting, then evaporated under nitrogen gas. Permethylated N-glycans were dissolved in 20 μl of 50% (v/v) methanol and co-crystallized with 2,5-dihydroxybenzoic acid matrix (10 mg/ml in 70% acetonitrile, 0.1% trifluoroacetic acid, and 2.5 mM sodium acetate). MALDI-TOF analysis was performed in positive reflector mode, acquiring 2,000 shots per spot in the m/z 1,000-7,000 mass range.

The resulting glycoprofile data was reanalyzed using GlyCompare^35^ to quantify all glycan intermediates (glyco-motifs) for each sample. Hierarchical clustering analysis was performed using the ward.D2 agglomeration method to identify mutants with similar glycan biosynthesis patterns. Prior to clustering, missing values in the glyco-motif abundance matrix were set to zero. The dendrogram was cut to identify three distinct clusters (k=3) representing different levels of glycosylation modification. For visualization, a symmetric color palette was generated using RColorBrewer’s RdBu palette, and the clustering results were displayed using annHeatmap2 with both row and column dendrograms. Nine major glyco-motif clusters (GM 01-09) were identified through column clustering, with each cluster annotated based on representative glycan topology shared by >60% of glyco-motif members within that cluster. The robustness of the clustering was assessed by comparing the branching patterns in the dendrograms.

### RNA sample preparation and sequencing

WT Rit, 6xKO Rit, 11xKO Rit cells (5 × 106) sampled from the DASGIP bioreactor on days 4, 6, and 8 were resuspended in RLT buffer (Qiagen, Hilden, Germany) containing 40-mM DTT. After storage at −80 °C, RNA was extracted using the RNeasy kit (Qiagen) followed by on-column DNAse digestion. RNA was eluted in 40 µl nuclease-free water, concentration was measured by Qubit (Thermo Fisher scientific) and the purity was checked on a Fragment Analyzer (Advanced Analytical). Complementary DNA synthesis was obtained from 1 μg of RNA using the High capacity cDNA RT kit (Thermo Fisher scientific). Samples were diluted to 60 ng µL−1 in 50 µL and library preparation was performed with the TruSeq Stranded mRNA Library Prep Kit (Illumina, San Diego, CA, USA). Final RNA libraries were first quantified by Qubit and the size was checked on a Fragment Analyzer. Libraries were normalized to 10 nM and pooled and the final pool of libraries was run on the NextSeq platform with high output flow cell configuration (NextSeq® 500/550 High Output Kit v2, 300 cycles, Illumina). Raw data are deposited at the Gene Expression Omnibus and Short Read Archive with accession number.

### RNA-Seq quantification and differential gene expression

RNA-Seq quality was assessed using FastQC. Adapter sequences and low quality bases were trimmed using Trimmomatic^81^. Sequence alignment was performed using STAR^82^ against the CHO genome (GCF_000419365.1_C_griseus_v1.0) with the default parameters. The expression of each gene was quantified using HTSeq^83^.

Gene counts were normalized using variance stabilizing transformation. To facilitate cross-species comparison, CHO genes were mapped to their mouse orthologs using a curated orthology mapping table. Gene symbols were obtained by mapping the mouse Entrez IDs to official gene symbols using the org.Hs.eg.db annotation package. For downstream analysis, we focused on glycosylation-related genes, including glycosyltransferases, glycan-binding proteins, and other glycosylation-related enzymes based on curated annotation lists.

Differential expression analysis was performed using DESeq2^84^ to compare each glycoengineered cell line to the wild-type control. For each comparison, we generated log2 fold changes and corresponding p-values. A gene was considered differentially expressed if it showed an absolute log2 fold change > 1.5 and adjusted p-value < 0.05. The results were compiled into a comprehensive differential expression table containing both the log2 fold changes and p-values for all genes across different glycoengineering conditions. For visualization and interpretation, differentially expressed genes were grouped by glycosyltransferase families (Mgat, B4galt, St3gal, and B3gnt) and their biological functions. Expression changes were visualized using hierarchical clustering with Ward’s method (ward.D2).

### Cluster comparison analysis

To assess the relationship between glycomotif patterns and gene expression, we performed a quantitative comparison of the hierarchical clustering results using the dendextend R package. The dendrogram structures were first aligned by identifying overlapping samples between datasets using tree intersection methods. The dendrograms were then optimized through a two-step process: an initial random search algorithm with 2,000 iterations, followed by stepwise rotation optimization to minimize dendrogram entanglement.

The similarity between clustering patterns was evaluated using the Fowlkes-Mallows (FM) index, which quantifies the agreement between two hierarchical clusterings. Statistical significance was assessed through permutation testing with 100,000 iterations. For each iteration, cluster labels were randomly permuted while maintaining the dendrogram structure, and the FM index was recalculated using the FM_index_R function. The empirical p-value was determined by comparing the observed FM index to the distribution of permuted values, with p < 1e-5 considered significant. The entanglement metric was also calculated to quantify the amount of crossing between dendrograms, with lower values indicating better agreement between the clustering patterns.

### Bioprofile FLEX Analyzer and data processing

The concentrations of glucose, lactate, ammonium (NH4^+^), and glutamine in spent media were measured using the BioProfile 400 (Nova Biomedical). Also measured key cellular phenotypes including cell size, growth rate, and viability.

Cell culture measurements were processed to calculate key metabolic and growth parameters. The exponential growth rate (r) was calculated using the equation N(t) = N(0)e^(rt), where N(0) and N(t) represent the viable cell density (VCD) at the initial and 48-hour timepoints, respectively. The cell population doubling time was derived as ln(2)/r.

Metabolic uptake and secretion rates were calculated for glutamine, glutamate, glucose, lactate, NH4^+^, Na^+^, K^+^, and osmolality. The specific rates were determined using the equation (C2-C1)/(0.5(t2-t1)(VCD1+VCD2)), where C2 and C1 are the metabolite concentrations at 48 hours and 0 hours, t2-t1 is the time interval (48 hours), and VCD1 and VCD2 are the viable cell densities at the respective timepoints.

Fold changes between 0 and 48 hours were calculated for cell size, VCD, and viability as the ratio of final to initial values. For baseline media composition, average values of initial metabolite concentrations were used across all samples to establish consistent reference conditions.

Data was first visualized using hierarchical clustering with Ward’s method (ward.D2). Then, for phenotype visualization, measurements (cell size, VCD, viability, and growth rate) were normalized by subtracting wild-type values. The normalized data was visualized using boxplots with individual data points overlaid using ggplot2. Samples were grouped by glycosyltransferase family (WT, Mgat, B4gal, ST3gal), with other perturbations classified as "Other". For each phenotype, the comparison between glycosyltransferase families was visualized using position-adjusted boxplots.

### Similarity Network Fusion Analysis

To integrate multiple-omics data types and identify patterns in cellular responses to glycoengineering, we employed Similarity Network Fusion (SNF)^36^ to construct and analyze patient-similarity networks. SNF enables integration of phenotypic and gene expression data by constructing a network for each data type and then fusing them into one comprehensive similarity network that captures both shared and complementary information.

We first constructed a sample similarity network represented as a graph G = (V,E), where vertices V correspond to the cell line samples and edges E are weighted by pairwise sample similarities. Edge weights were computed using a scaled exponential similarity kernel. We employed normalized mutual information (NMI) to validate the compatibility of the integrated data sources. All SNF analyses were performed using the SNFtool R package with default parameters unless otherwise specified.

### Gene Set Variation Analysis

Prior to regression, pathway activity scores were computed using Gene Set Variation Analysis (GSVA)^85^. RNA-Seq count data were normalized via variance-stabilizing transformation and mapped to mouse orthologs. Reactome pathways^37^ were selected for their relevance to glycosylation, metabolism, and stress responses. GSVA enrichment scores quantified the relative activity of each pathway across samples, enabling systematic comparison of metabolic and functional shifts associated with glycoengineering.

### Ordinal regression analysis

Ordinal regression was performed using cumulative link models (CLMs)^86^ to analyze ordered categorical outcomes. The analysis was implemented in R using the ordinal package’s clm function, which estimates threshold parameters and predictor effects simultaneously while maintaining the ordered nature of response categories. Pathway activity scores, derived from Gene Set Variation Analysis (GSVA), were modeled as continuous responses, with the number of targeted glycosyltransferase genes (e.g., St3gal, Mgat, or B4galt family) knockouts as an ordinal predictor.

The CLM framework assumes proportional odds, where the effect of increasing knockouts on pathway activity is consistent across thresholds. Coefficients (ORscore) were interpreted as log-odds ratios, where positive values indicated pathways upregulated with increasing knockouts, and negative values indicated downregulation. ORPvalue is the statistical significance of the ORscore, derived from the CLM. Significance thresholds were set at an absolute log2 fold change >1.5 and adjusted p-value <0.05 after Benjamini-Hochberg correction for multiple testing.

Pathways with significant ordinal trends (p < 0.05) were visualized as heatmaps (Figure 6a, S2a, S3a). Top 3 up- and down-regulated pathways were visualized as box plots (Figure 6b, S2b, S3b).

### CHO-S derived recombinant protein purification and data collection

Herceptin, Rituximab and Enbrel were purified by protein A affinity chromatography. For each protein glycoform, a 1-mL MAbSelect Extra column (Cytiva) was equilibrated with 5 column volumes (CV) of 20 mM sodium phosphate, 0.15 M NaCl, pH 7.2. Next, 30-mL supernatant was loaded, the column was washed with 20 CV of 20 mM sodium phosphate, 0.15 M NaCl, pH 7.2, and the protein was eluted using 0.1 M citrate, pH 3.0. The elution fractions (0.5 mL) were collected in deep-well plates containing 100 µL of 1 M Tris at pH 9 per well.

Alpha-1-antitrypsin, butyrylcholinesterase, protein Z, serpin A5, serpin A10, serpin C1, and EPO, all C-terminally tagged with the HPC4 tag (amino acids EDQVDPRLIDGK), were purified over a 1-mL column of anti-protein C affinity matrix according to the manufacturer’s protocol (Roche, cat. no. 11815024001). 1 mM CaCl_2_ was added to the supernatant, equilibration buffer, and wash buffer, respectively. The proteins were eluted in 0.5 mL fractions using 5 mM EDTA in the elution buffer.

For all proteins, RNA-Seq quality was assessed using FastQC. For details, refer to the RNA-Seq quantification section above. Following RNA-seq processing, Gene Ontology (GO)^87^ term analysis was performed to annotate biological pathways. GSVA was then applied to quantify pathway activity, generating enrichment scores for each sample. Samples were stratified into two groups based on knockout status (0 vs. 3) to compare pathway dysregulation. Differential pathway activity between groups was assessed using the Wilcoxon rank-sum test, and pathways with statistically significant differences (p < 0.05) were retained. From these, DNA damage repair-related pathways were selectively identified based on functional relevance. To visualize the most pronounced changes, the top three pathways exhibiting the strongest upregulation and downregulation (ranked by effect size) were plotted as boxplots, illustrating the distribution of GSVA scores across knockout conditions.

### Statistical Analyses

Statistical analyses were performed using R (version 4.1.0), and data visualization was carried out using custom R scripts incorporating the ggplot2 and related packages. Principal Component Analysis (PCA) was performed using the PCA function from FactoMineR package.

## Data Availability

The authors declare that all other data supporting the findings of this study are available within the paper and its supplementary information files.

## Code Availability

The computing environment was standardized across all experiments, running on Apple M2 chip.

## Acknowledgments

This work was supported by generous funding from NIGMS (R35 GM119850) and the Novo Nordisk Foundation (NNF20SA0066621). The authors also wish to thank Zhang Yang, Hiren Joshi, and Henrik Clausen for sharing cell lines used in this study.

