## Supplementary figures for "Delineating the Transcriptional and Phenotypic Impact from Biotherapeutic Glycoengineering"

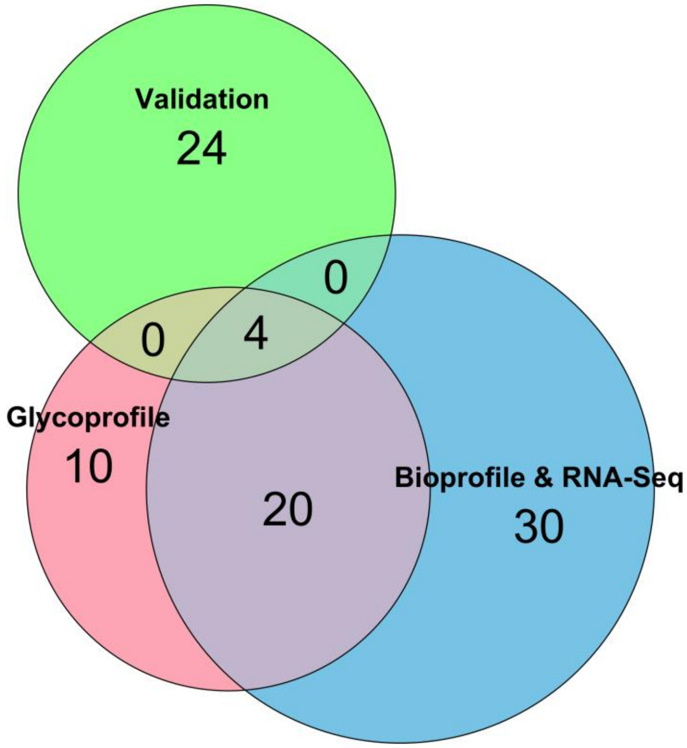


**Figure S1. Venn diagram illustrating overlap and uniqueness among Validation, Glycoprofile, and Bioprofile & RNA-Seq datasets.** The Glycoprofile dataset comprises 34 genotypes, of which 24 overlap with the Bioprofile & RNA-Seq dataset. The Bioprofile & RNA-Seq dataset contains 30 unique genotypes not shared with other datasets. The Validation dataset (RNA-Seq) includes 24 unique genotypes.

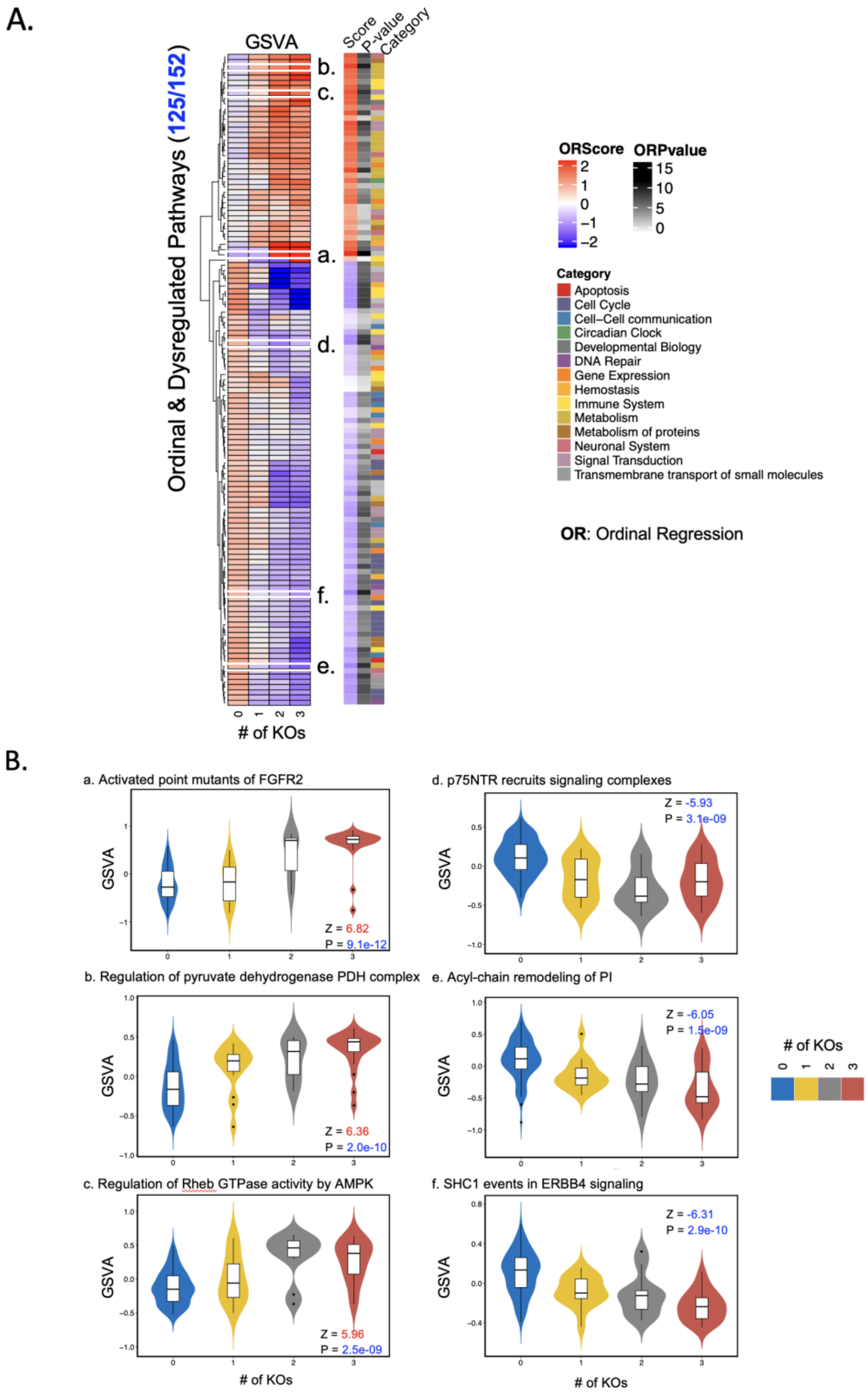


**Figure S2. Ordinal regression results of GSVA of pathways in the Mgat-family glycoengineered strains.** (A) The heatmap of statistically significant dysregulated pathways with an ordinal trend of GT knockouts. (B) Statistically significant dysregulated pathways.

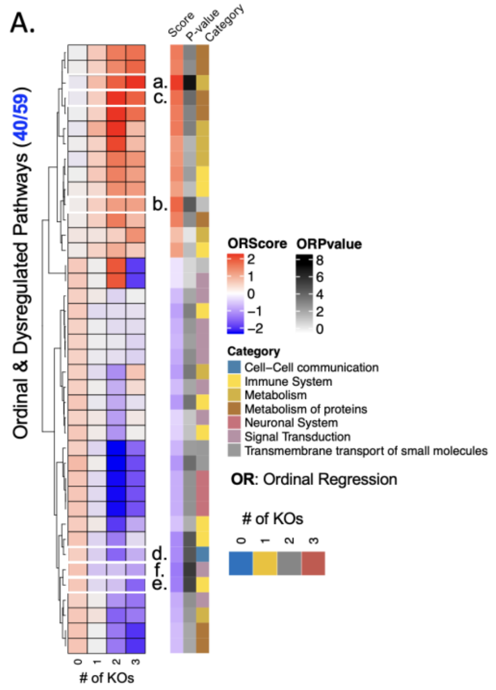


**Figure S3. Ordinal regression results of GSVA of pathways in the B4galt-family glycoengineered strains.** (A) The heatmap of statistically significant dysregulated pathways with an ordinal trend of GT knockouts. (B) Statistically significant dysregulated pathways.

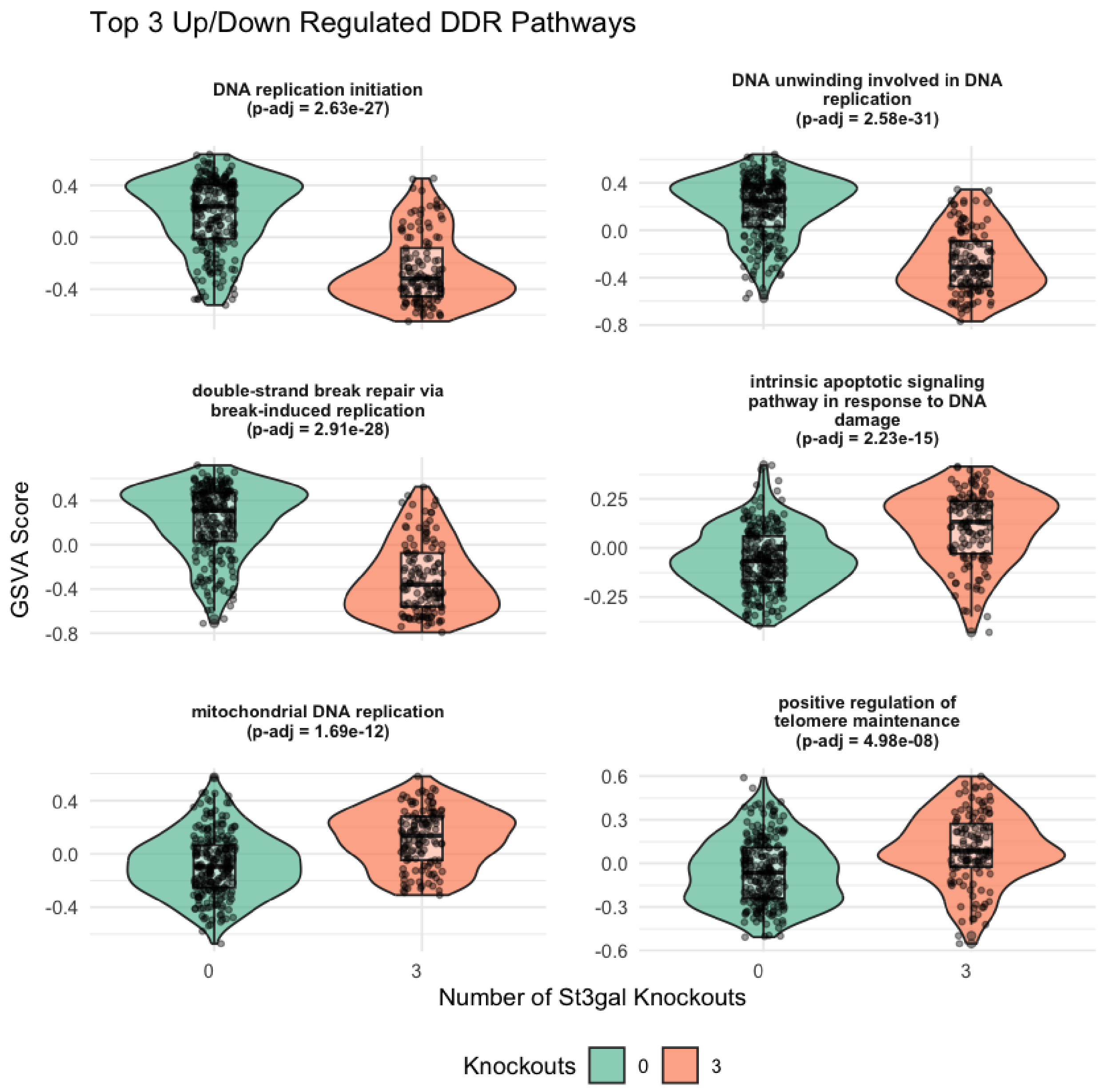


**Figure S4. Validation results of GSVA of pathways in the St3gal-family glycoengineered strains.** Statistically significant dysregulated DNA-damage-repair related pathways.
